# HepaRG cells undergo increased levels of post-differentiation patterning in physiologic conditions when maintained as 3D cultures in paper-based scaffolds

**DOI:** 10.1101/2023.01.16.524330

**Authors:** Thomas J. Diprospero, Lauren G. Brown, Trevor D. Fachko, Matthew R. Lockett

**Affiliations:** Department of Chemistry, University of North Carolina at Chapel Hill, Kenan and Caudill Laboratories, Chapel Hill, NC 27599-3290, United States; Lineberger Comprehensive Cancer Center, School of Medicine, University of North Carolina at Chapel Hill, Chapel Hill, NC 37599-7295, United States

**Keywords:** Hepatocyte, Cytochrome P450 (CYP), Monolayer, Oxygen, Wnt/β-Catenin

## Abstract

Monolayer cultures of hepatocytes lack many aspects of the liver sinusoid, including a tissue-level organization that results from extracellular matrix interactions and gradients of soluble molecules that span from the portal triad to the central vein. We measured the activity and transcript levels of drug-metabolizing enzymes in HepaRG cells maintained in three different culture configurations: as monolayers, seeded onto paper scaffolds that were pre-loaded with a collagen matrix, and when seeded directly into the paper scaffolds as a cell-laden gel. Drug metabolism was significantly decreased in the presence of the paper scaffolds compared to monolayer configurations when cells were exposed to standard culture conditions. Despite this decreased function, transcript levels suggest the cells undergo increased polarization and adopt a biliary-like character in the paper scaffolds, including the increased expression of transporter proteins (e.g., *ABCB11* and *SLOC1B1*) and the *KRT19* cholangiocyte marker. When exposed to representative periportal or perivenous culture conditions, we observed in vivo zonal-like patterns, including increased cytochrome P450 (CYP) activity and transcript levels in the perivenous condition. This increased CYP activity is more pronounced in the laden configuration, supporting the need to include multiple aspects of the liver microenvironment to observe the post-differentiation processing of hepatocytes.

**TOC Figure:** 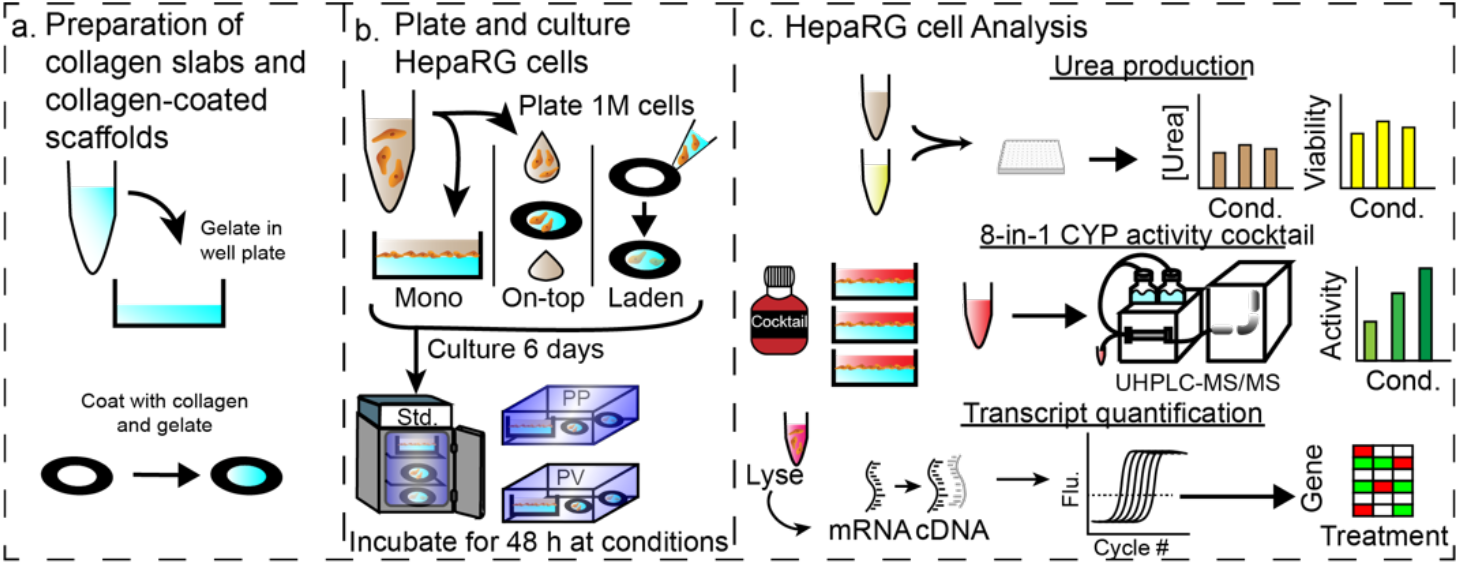

## Introduction

Drug-induced liver injury and hepatotoxicity are key contributors to late-stage drug failure.(Kenna and Uetrecht 2018; Parasrampuria et al. 2018) These failures arise, in part, to the limited predictive power of the cell-based assays and animal models currently relied upon in the drug discovery pipeline, allowing potentially toxic drugs to reach the patients. Cell-based assays rely on monolayers of primary human hepatocytes (PHHs), which readily dedifferentiate in this culture format.(Godoy et al. 2009) Monolayer formats also expose cells to a uniform culture condition, instead of the gradients of oxygen, nutrients, and other soluble factors that span the liver sinusoid. These gradients of soluble molecules result in liver zonation,(Usta et al. 2015) the spatially regulated distribution of liver-specific function along each sinusoid.(Kietzmann 2019) The oxygen gradient is well characterized and thought to be a master regulator of hepatic function, including the expression of drug-metabolizing enzymes.(Kietzmann 2017; Matz-Soja et al. 2013) Oxygen partial pressures range from pO_2_ = 11% in the periportal (PP) region to pO_2_ = 3% in the perivenous (PV) region. We, and others, showed that placing monolayers of hepatocyte-like cell lines under decreasing oxygen tensions resulted in increased cytochrome P450 (CYP) activities, aligning with zonal localization found in vivo.(DiProspero et al. 2022; DiProspero et al. 2021; van Wenum et al. 2018) Signaling molecules and morphogens secreted by non-parenchymal cells also regulate key hepatic functions.(Gebhardt and Hovhannisyan 2010) The Wnt ligand, which is localized in the PV region, promotes the expression of glutamine synthase.(Benhamouche et al. 2006) We previously evaluated the impact of Wnt3a, R-spondin, and noggin (WRN) molecules on the expression and activity of drug-metabolizing enzymes in monolayers of HepaRG cells.(Kyffin et al. 2018) Under standard culture conditions and atmospheric oxygen partial pressures, the WRN molecules significantly downregulated CYP activity. However, under representative PP or PV oxygen partial pressures, the WRN molecules enhanced the oxygen-regulated patterning of CYP activity.

In addition to extracellular gradients of molecular modulators, the sinusoid has a distinct architecture where hepatocytes are maintained in an extracellular matrix (ECM)-rich environment. The addition of ECM in the form of a collagen sandwich was an early advance in hepatocyte culture,(Gaskell et al. 2016; Zhou et al. 2019) prolonging the differentiated state of PHHs in vitro while retaining standard medium and culture conditions. This sandwich structure maintained polarized primary mouse hepatocytes and functional bile canaliculi for twice as long as monolayers on collagen slabs.(Choi and Choi 2013) Another advance in hepatocyte cultures is the generation of metabolically competent, hepatocyte-like cell lines. The HepaRG cells used in this work are human bipotent progenitors, which differentiate into cholangiocyte-like and hepatocyte-like cells.(Parent et al. 2004) The hepatocyte-like cells have drug-metabolizing profiles similar to PHHs, expressing phase I, II, and III proteins.(Wang et al. 2015) The health and drug-metabolizing activity of differentiated HepaRG cells is culture format-dependent. HepaRG spheroids secreted higher amounts of albumin (2-fold) and had increased CYP3A4 activity (1.5-fold) compared to monolayer formats.(Higuchi et al. 2016) The improved predictiveness of spheroids is attributed to increased cell-cell interactions and the formation of mass transport-limited gradients.(Bell et al. 2016; Cox et al. 2020) A limitation of spheroids is the inability to determine which microenvironment components affect cellular function in a spatially resolved manner. Microfabricated devices have incorporated sinusoid-like architectures with experimentally defined oxygen and nutrient gradients imposed across mono- and co-culture formats.(Dalsbecker et al. 2022; Starokozhko and Groothuis 2017) Kang developed a microfluidic system that measures glucose metabolism in hepatocytes along physiologically relevant gradients of either oxygen(Kang et al. 2020) or hormones.(Kang et al. 2018)

We recently showed that a combination of physiologic oxygen tensions and signaling molecules promoted zonal-like post-differentiation of drug-metabolizing enzyme activity in monolayer cultures of HepaRG cells.(DiProspero et al. 2022) To expand on these findings and determine if tissue-like structures can improve zonation, we incorporated paper scaffolds seeded with collagen. The first example of paper as a readily accessible scaffold for supporting cell-laden hydrogels was presented by Whitesides.(Derda et al. 2009) In the past decade, paper scaffolds have been used to assemble tissue-like structures for fundamental biological studies and translational applications in tissue repair and engineering.(Cramer et al. 2019) Paper scaffolds can support the prolonged culture of hepatocytes,(Agarwal et al. 2020a; Wang et al. 2018) and recently were highlighted as a promising material for engineering liver tissues.(Agarwal et al. 2020b)

To characterize the effect of the paper scaffolds on HepaRG cellular function, we quantified urea secretion, the expression and activity of drug-metabolizing enzymes, and the expression of polarization and cholangiocyte markers in cells maintained in the culture configurations shown in **Figure 1**. The progression from the monolayer to laden configuration increased the dimensionality of the culture while also assessing if the paper scaffold affected cell health or function. The comparison of the monolayer and on-top configuration evaluated the placement of the culture, as the paper scaffolds remained at the air-water interface throughout the experiment. We assessed the cells under standard and physiologically relevant culture conditions. Standard culture conditions of pO_2_ = 20% and basal medium served as the baseline for comparison. To model the PP region, we maintained the cells at pO_2_ = 11% O_2_ and in basal medium; for the PV region, we maintained the cells at pO_2_ = 5% and in medium containing the WRN molecules.

**Figure 1.**
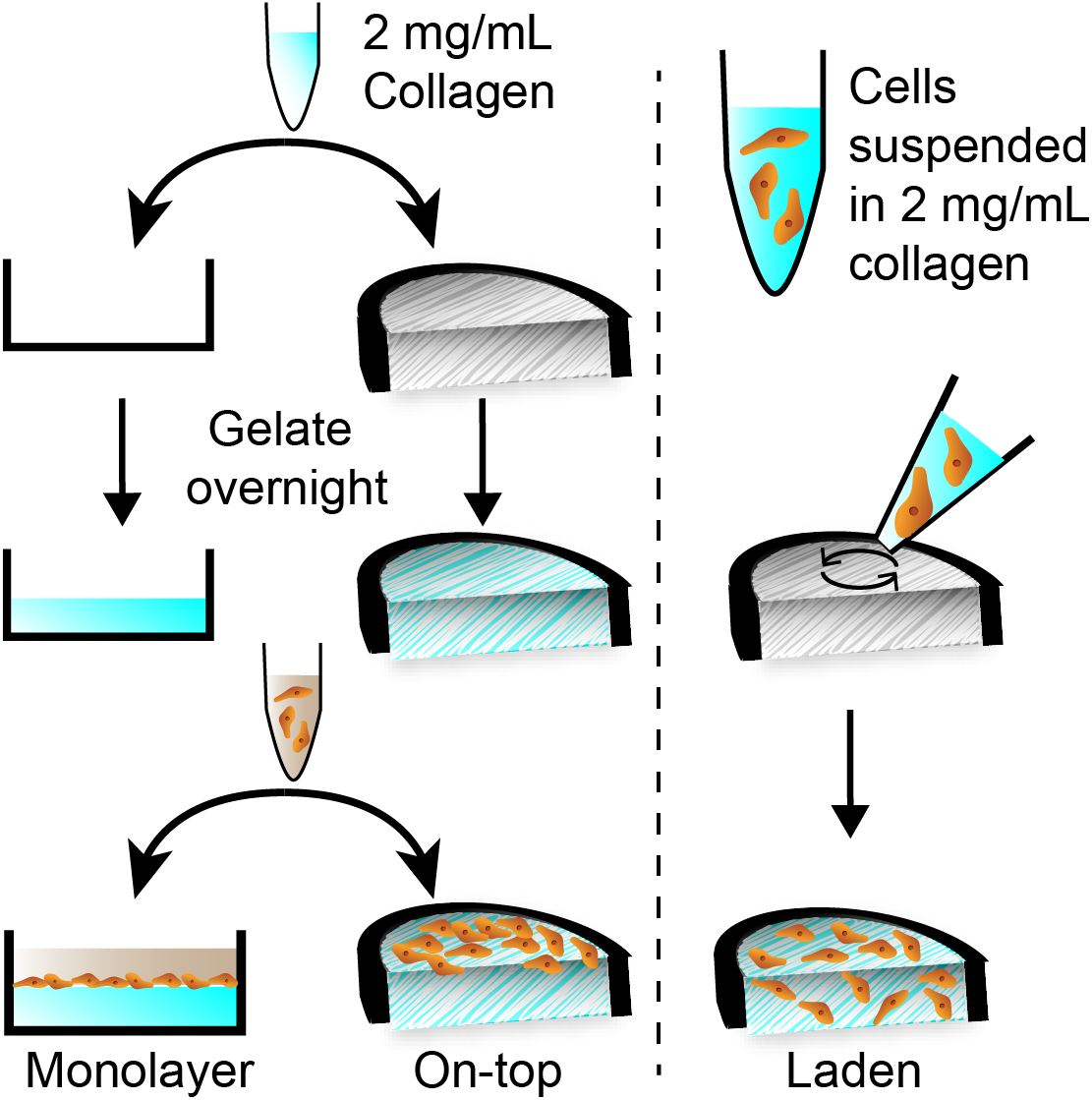
Overview of the three culture configurations evaluated in this work, each containing collagen I (2 mg/mL). In the monolayer configuration, the collagen was deposited onto the bottom of commercial 12-well plates and gelated overnight at 37 °C. In the on-top configuration, the collagen was deposited into the paper scaffolds and gelated overnight at 37 °C. The HepaRG cells (1.0×10^6^) were deposited onto the gelated collagen in both the monolayer and on-top configurations. In the laden format, 1.0×10^6^ HepaRG cells were suspended in collagen I, deposited into the paper scaffold, and gelated overnight at 37 °C.

Urea synthesis and the number of viable cells were unchanged between the three configurations under standard culture conditions, indicating the scaffolds did not inhibit cellular health. The monolayer HepaRG culture configuration under standard conditions had the highest CYP activity. With the inclusion of representative PP and PV culture conditions, we observed synergistic effects between the paper cultures and physiologic conditions. The paper scaffolds enhance the HepaRG response to the PP and PV environments, resulting in further increases in CYP activity and transcriptional regulation than when maintained as monolayers. These results highlight the need for increased context when assessing hepatocytes, including appropriate oxygenation levels and a 3D culture environment.

## Materials and methods

### Chemicals

All chemicals and reagents were used as received unless otherwise specified. Chlorzoxazone, dextromethorphan (hydrobromide hydrate), (S)-mephenytoin, midazolam, testosterone, and acetaminophen-d4 were purchased from Cayman Chemical Company. Collagen I (rat tail) was purchased from Enzo Life Sciences. Phenacetin, dextrorphan-d3, bupropion hydrochloride, and sodium hydroxide (NaOH) were purchased from Millipore Sigma. Dimethyl sulfoxide (DMSO) and 7-hydroxycoumarin were purchased from Fisher Scientific. Hydroxybupropion-d_6_ was purchased from Cambridge Isotope Laboratories. 6-hydroxychlorzoxazone-d_2_ was purchased from Clearsynth. 7-hydroxycoumarin-d5-sulfate and 7-hydroxycoumarin-^13^C_6_-glucuronide were purchased from Toronto Research Chemicals. 4-hydroxymephenytoin-d_3_ was purchased from MuseChem. Hydroxymidazolam-^13^C_3_ and hydroxytestosterone-d_7_ were purchased from Corning.

### Preparation of the paper scaffolds

Paper scaffold preparation and sterilization were detailed previously.(Kenney et al. 2019; Lloyd et al. 2017) Briefly, sheets of Whatman 105 lens paper were patterned with wax borders using a Xerox ColorQube 8570 printer. Each scaffold was 18 mm in diameter and fit directly into the well of a standard 6-well plate. The scaffolds contained a 4 mm wax border that defined the cell culture region. The wax border also ensured the scaffolds remained at the air-medium interface during culture. Prior to use, the paper scaffolds were sterilized under UV light for one hour. **Figure S1** contains photographs and detailed schematics of the paper scaffolds.

### LWRN cell culture and conditioned media collection

The L- and L-WRN cell lines were purchased from ATCC. Both cell lines were maintained as monolayers at 20% O_2_, 37 °C, and 5% CO_2_ in Dulbecco’s Modified Eagle’s Medium (DMEM) supplemented with 10% fetal bovine serum (FBS), 0.5 mg/mL G-418, and 0.5 mg/mL hygromycin B. Cell culture medium and supplements were purchased from Gibco. The culture medium was exchanged every 2-3 days, and the cells were passed at 80% confluency with TrypLE using standard procedures. The conditioned medium was collected with the ATCC-recommended protocol. Once collected, the medium was sterile filtered (0.22 μm) and stored at −80°C.

### HepaRG cell culture

Differentiated NoSpin HepaRG cryopreserved cells, medium, and supplements were purchased from Lonza Bioscience. 1.0×10^6^ HepaRG cells were deposited onto collagen slabs at the bottom of a well plate, deposited onto sheets of collagen-laden paper, or suspended in collagen and seeded into the paper sheets. **Figure 1** summarizes these culture configurations and their preparation. Once deposited, the HepaRG cells were maintained at 37 °C, 20% O_2_, and 5% CO_2_ unless otherwise noted. For the first 24 h, the cells were maintained with HepaRG medium supplemented with a basal supplement, a thawing and plating supplement, and 1% penicillin-streptomycin (PenStrep, Gibco). After a medium exchange, the cells were maintained in HepaRG medium containing a basal medium supplement, a maintenance and metabolism supplement, and 1% PenStrep. The maintenance medium was exchanged every 2 d. On day 6, the cells were exposed to the pO_2_ values and culture medium combinations detailed in the Results section. Oxygen tensions were regulated in a custom-built hypoxia chamber, as detailed previously.(DiProspero et al. 2021)

The collagen supports were prepared by neutralizing an acidified suspension of collagen I with NaOH and phosphate-buffered saline (1X PBS) to a pH = 7.2. The final density of the collagen suspension was adjusted to 2 mg/mL with RO water before use. Commercial 12-well plates were coated with the neutralized collagen solution, incubated overnight at 37°C, and washed once with 1X PBS prior to cell deposition. Paper scaffolds were loaded with the neutralized collagen solution (25 μL), incubated overnight at 37°C in 1X PBS, and washed once with 1X PBS prior to cell deposition. Cell-laden paper scaffolds were seeded with 12.5 μL of collagen containing 80,000 cells/μL suspension of HepaRG cells.

### Evaluation of metabolic enzyme activity

CYP activity was quantified using an LC-MS/MS method we adapted from a previous protocol.(DiProspero et al. 2022; Li et al. 2015) The HepaRG cells were washed with 1X PBS and incubated for 2 h at 37 °C in HepaRG medium containing a basal medium supplement and eight different enzyme substrates (**Table S1**). Stock solutions of each substrate were prepared in DMSO at 1000X the working concentration and added to the HepaRG medium directly before use. After 2h, the cell culture medium was mixed with ice-cold acetonitrile (−20 °C, Optima) containing isotopically labeled standards of each enzyme product at a 1:10 (v/v) ratio. Precipitated protein was removed with centrifugation (12,000 xg, 15 min), the supernatant collected, and then concentrated in vacuo. The residual solids were resuspended in 100 μL of HPLC-grade water (Optima).

Samples were separated on an Acquity UPLC (Waters) equipped with a BEH C18 column (2.1 x 50 mm, 1.7 μm) using a binary solvent system of (A) 0.1% formic acid (v/v) in water and (B) 0.1% formic acid (v/v) in acetonitrile. The total run time of each separation was 9 min, using the following gradient profile at a 0.3 mL/min flow rate: 10% B for 1 min; a linear gradient to 70% B over 5 min; 95% B for 1 min; 10% B for 2 min to re-equilibrate the column. Enzyme products were quantified with selected reaction monitoring (SRM) on a Thermo TSQ Vantage triple quadrupole mass spectrometer (MS). The MS source parameters were a heated electrospray ionization probe, 300 °C; capillary temperature, 300 °C; spray voltage, 4.8 kV; vaporization temperature, 300 °C; sheath gas flow, 50; auxiliary gas flow, 15; S-lens RF amplitude, 120 V. Nitrogen was used for the sheath and auxiliary gases. Argon was used as the collision gas. Two transitions of each product and internal standard were monitored except for 4’-hydroxymephenytoin-d3 where only a single transition could be monitored due to instrument limitations. The optimized declustering voltage and collision energies of each product and internal standard are listed in **Table S1**. Data were collected and processed with the Xcalibur software package. A matrix control was included in each run to ensure no interferents co-eluted.

Quality control samples containing 10 μM of each analyte were analyzed after every 10 injections to monitor instrument drift. The analyte-to-internal standard (IS) ratio of each enzyme product was averaged across three separately prepared cell cultures.

### Urea production

Urea concentration was determined with the Quantichrom Urea Assay Kit (BioAssay Systems), according to the manufacturer’s protocol. Prior to analysis, the cells were washed with 1X PBS and placed in fresh culture medium. An aliquot of the medium was measured (absorbance at 430 nm) on a SpectraMax i3x Microplate Reader, and concentrations were determined from a calibration curve.

### Viable cell count

Cells were washed with 1X PBS and incubated in a tri-color stain of Hoechst 33342 (10 μg/mL), calcein-AM (2 μg/mL), and PI (3 μg/mL) in 1X PBS for 10 min at room temperature. Prior to imaging, the cells were washed three times with 1X PBS. Images were collected on a Nikon TE-2000i inverted microscope with a CoolSNAP DYNO CCD camera. The fraction of viable cells was calculated with Eqn. 1, where the number of dead cells *(n_dead_)* was determined by counting PI-stained nuclei and the total number of cells *(n_all_)* from Hoechst-stained nuclei.

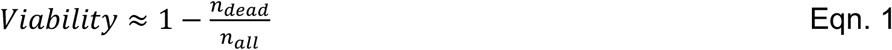

### Transcript Expression Quantification with RT-qPCR

The cell-containing paper scaffolds were washed with 1X PBS, placed in a 1.5 mL microcentrifuge tube containing 1.0 mL of TRIzol, vortexed for 30 s, and incubated at room temperature for 5 min with agitation. The RNA was collected using a TRIzol Plus purification kit (ThermoFisher). Reverse transcription was performed immediately after RNA isolation with a High-Capacity cDNA Reverse Transcription Kit (ThermoFisher) in an Eppendorf Master Cycler. Amplification reactions for qPCR were prepared in a 384-well plate with PowerUp SYBR Master Mix (ThermoFisher). Each gene was measured in triplicate on a QuantStudio 6 Flex Real-Time PCR system using the following program: 95 °C for 60 s, followed by 40 cycles of 95 °C for 2 s, and 60 °C for 30 s. **Table S2** lists the primer sequences, optimal primer concentrations, and reaction efficiencies (90-110%) of each gene of interest. Transcripts were quantified using the ΔΔCt method against *18sRNA*.(US Food and Drug Administration 2020)

### Statistical Analysis

All datasets are reported as the average and standard error of the mean (SEM) of at least four independent measurements prepared from at least two separate vials of HepaRG cells. Analyses were performed with GraphPad Prism 7 and the statistical tests detailed in the figure captions. An average fold-change of greater than two for qPCR datasets was considered significant. For CYP activity quantification, an average fold-change outside the window of 0.8 - 1.25 was considered significant with a *p*-value < 0.0001. These values were determined by analyzing consecutive injections of product standards (**Figure S2**). The remainder of the datasets collected were considered statistically significant for differences corresponding to a *p*-value of ≤ 0.05.

## Results

### HepaRG health and viability are unaffected by culture configuration

Previous characterization of cryo-preserved HepaRG monolayers found the basal-level activity of drug-metabolizing enzymes varied with time.(Jackson et al. 2016) **Figure 2** plots the drug-metabolizing enzyme activity of 1.0×10^6^ HepaRG cells in each culture configuration, maintained under standard culture conditions for 24 d. The onset of individual CYP activities varied, with the most significant increases in total CYP activity occurring on day 11 in the monolayer configuration and day 7 for both the on-top and laden configurations. Based on these datasets, we analyzed the HepaRG cells in all subsequent experiments 8 d after cell deposition.

**Figure. 2.**
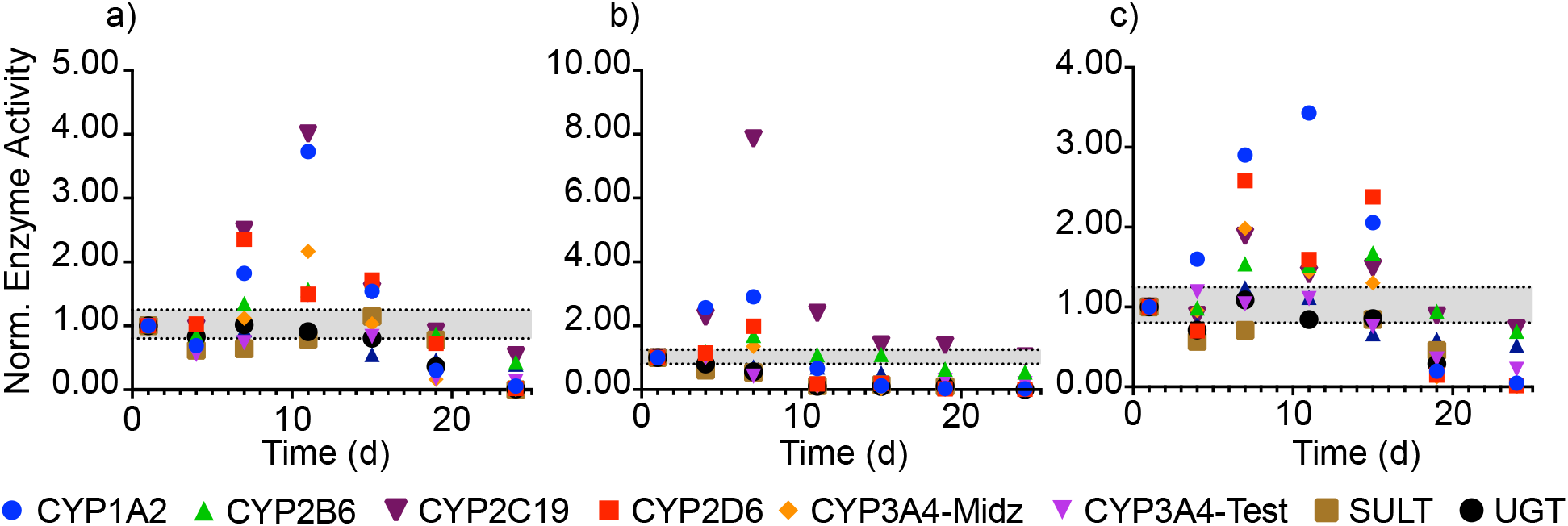
Drug-metabolizing enzyme activity of 1.0×10^6^ HepaRG cells maintained under standard culture conditions for 24 d after seeding in the a) monolayer, b) on-top, and c) laden culture configurations. Enzyme activity was determined with an LC-MS/MS method that quantified each enzyme product against its isotopically labeled internal standard. Values are the average of two separately prepared setups reported as a normalized activity to day 1. The dotted black lines represent a significantly different activity based on our LC-MS/MS method.

**Figure 3** shows representative micrographs of live HepaRG cells in each configuration stained with Hoechst 33342, PI, and calcein-AM dyes, 7 d after deposition. The percentage of viable cells in each culture configuration was above 80%, as determined from their nuclear stains. These values, along with a confirmation of viability from the calcein stains, suggest the time-dependent changes in drug-metabolizing enzyme activity were not due to cellular death. The cells are distributed evenly in a single focal plane in the monolayers and on-top configurations. In the laden configuration, the cells are distributed throughout the scaffold but preferentially aggregated in regions with a high density of overlapping cellulose fibers. The cells are likely not attaching directly to the fibers but rather to a difference in the collagen stiffness or density in those regions. The Supporting Information contains representative grey-scale images of each stain (**Figure S3**).

**Figure. 3.**
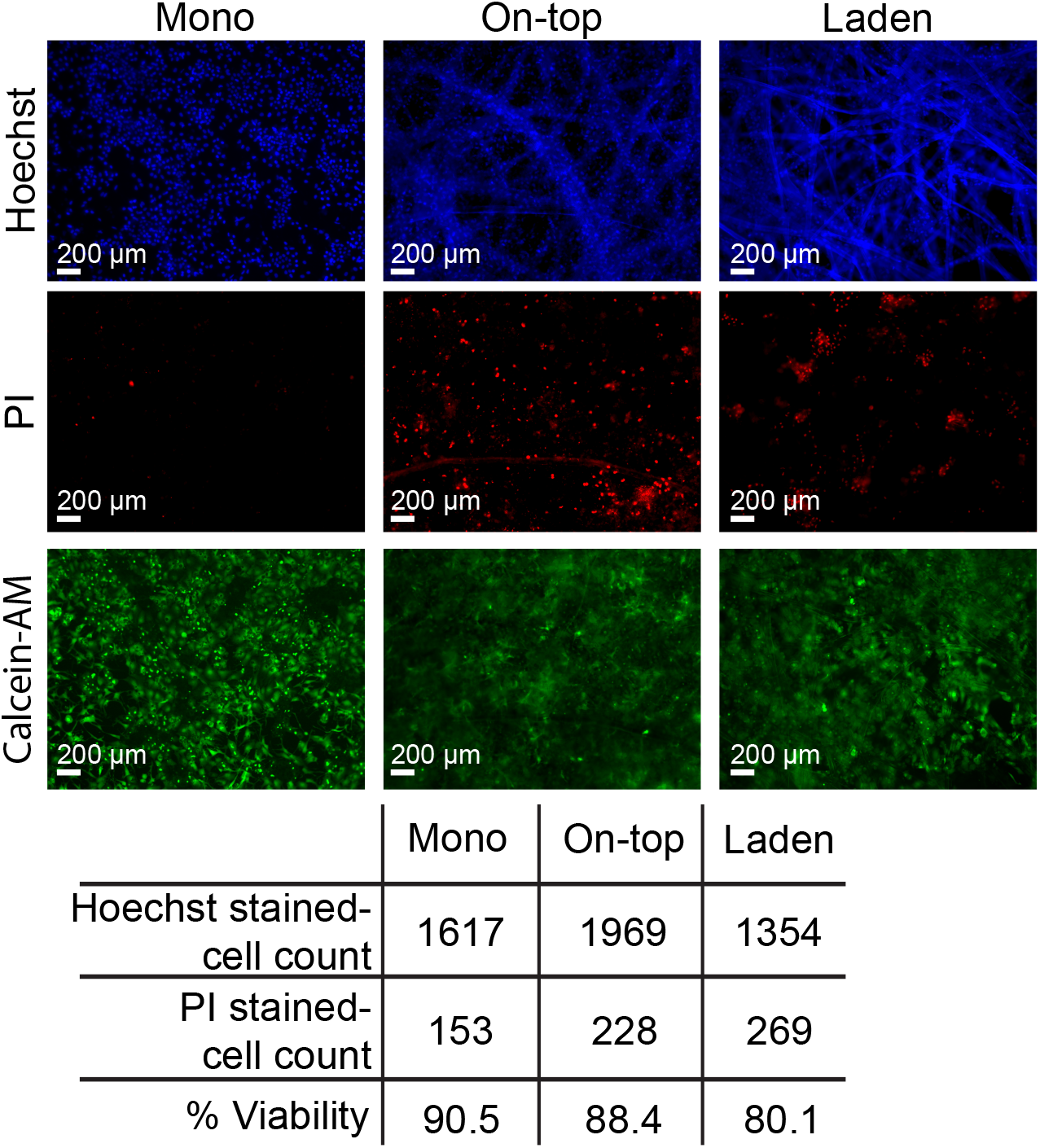
Representative images of 1.0×10^6^ HepaRG cells in each culture configuration, maintained under standard culture conditions, 7 d after seeding. The nuclei in each configuration were stained with Hoechst 33342 (total cell count) and PI (dead cell count). Cells with intact membranes were stained with calcein-AM. Percent viability was calculated using Eqn. 1 and represents the nuclei not stained with PI.

**Figure 4** plots the urea secretion from 1.0×10^6^ HepaRG in each culture configuration, maintained under standard culture conditions, 8 d after seeding. These values were not significantly different: 0.020 mg/mL/h/10^6^ cells in the monolayers, 0.010 mg/mL/h/10^6^ cells in the on-top configuration, and 0.011 mg/mL/h/10^6^ cells in the laden configuration. **Figure S4** shows urea secretion in each culture configuration was equivalent and unchanging over a 24-d period.

**Figure. 4.**
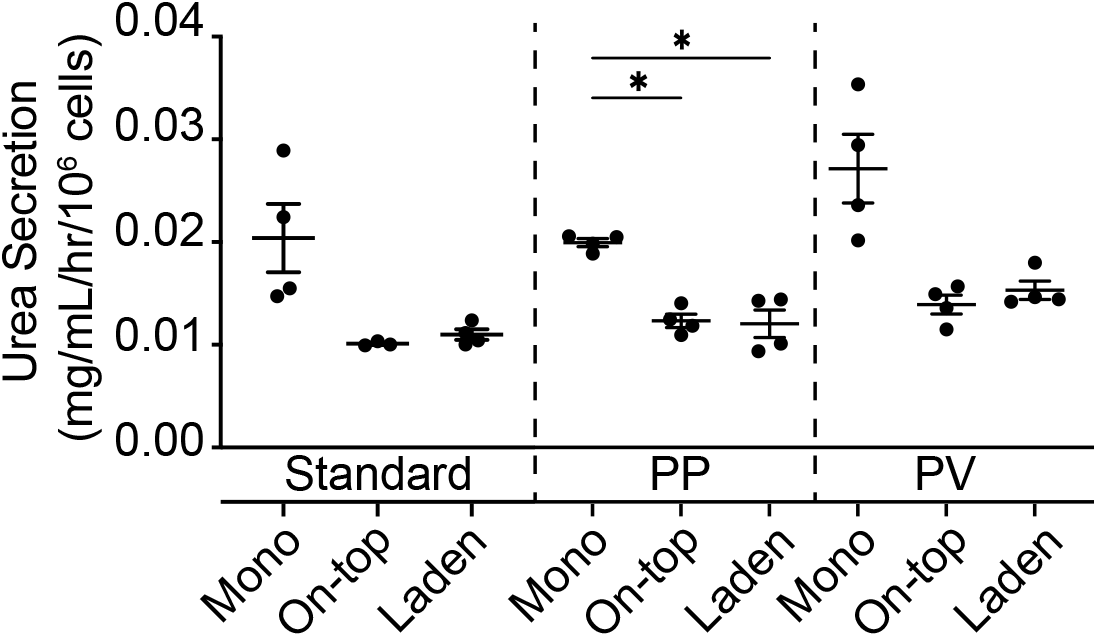
Urea secretion from 1.0×10^6^ HepaRG cells maintained (left) under standard culture conditions for 8 d, (middle) under standard culture conditions for 6 d followed by 2 d under PP conditions of pO_2_ = 11%, or (right) under standard culture conditions for 6 d followed by 2 d under PV conditions of pO_2_ = 5% pO_2_ in the presence of L-WRN conditioned medium. Datasets represent the mean and SEM of four replicate setups, prepared two cryovials of cells. Significance was determined with a one-way ANOVA, compared to the monolayer configuration under the same culture conditions.

### Basal drug-metabolizing enzyme activity under standard culture conditions decreased in the paper scaffolds

**Figure 5** plots the basal drug-metabolizing enzyme activity of the 1.0×10^6^ HepaRG cells 8 d after seeding, maintained under standard culture conditions. CYP1A2, 3A4, SULT, and UGT activity decreased significantly in the on-top configuration compared to the monolayer cultures. In the laden configuration, the average activity of CYP1A2, 2B6, 2C19, 2D6, 3A4, SULT, and UGT decreased significantly.

**Figure. 5.**
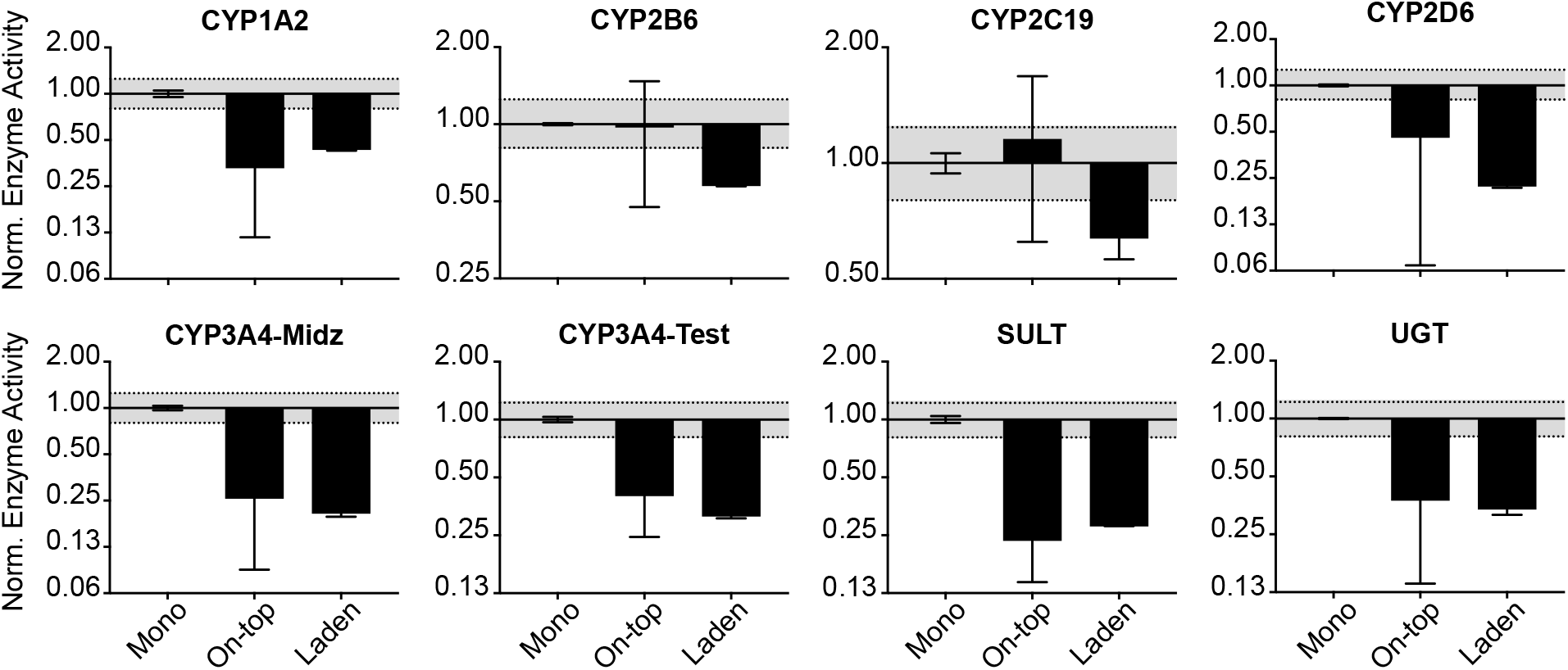
Basal drug-metabolizing enzyme activity of 1.0×10^6^ HepaRG cells, maintained under standard culture conditions, 8 d after seeding. Activity values were normalized to the monolayer configuration. Each bar represents the average and SEM of three separate setups, prepared from two cryovials of HepaRG cells. The dotted black lines represent measurable differences in activity, based on our LC-MS/MS method.

**Figure 6** displays the average transcript-level changes of the same HepaRG cells maintained under standard culture conditions, normalized to the monolayer configuration. *CYP2D6* and *3A4* levels were unchanged between the three configurations. *CYP1A2, 2B6, 2C19,* and *2E1* increased significantly in the on-top configuration. In the laden configuration, only *CYP1A2* and *UGT2B4* increased significantly. Increased *AHR* transcripts in the on-top (3.7-fold) and laden (4.2-fold) configurations correlate with increased *CYP1A2* transcripts.

**Figure. 6.**
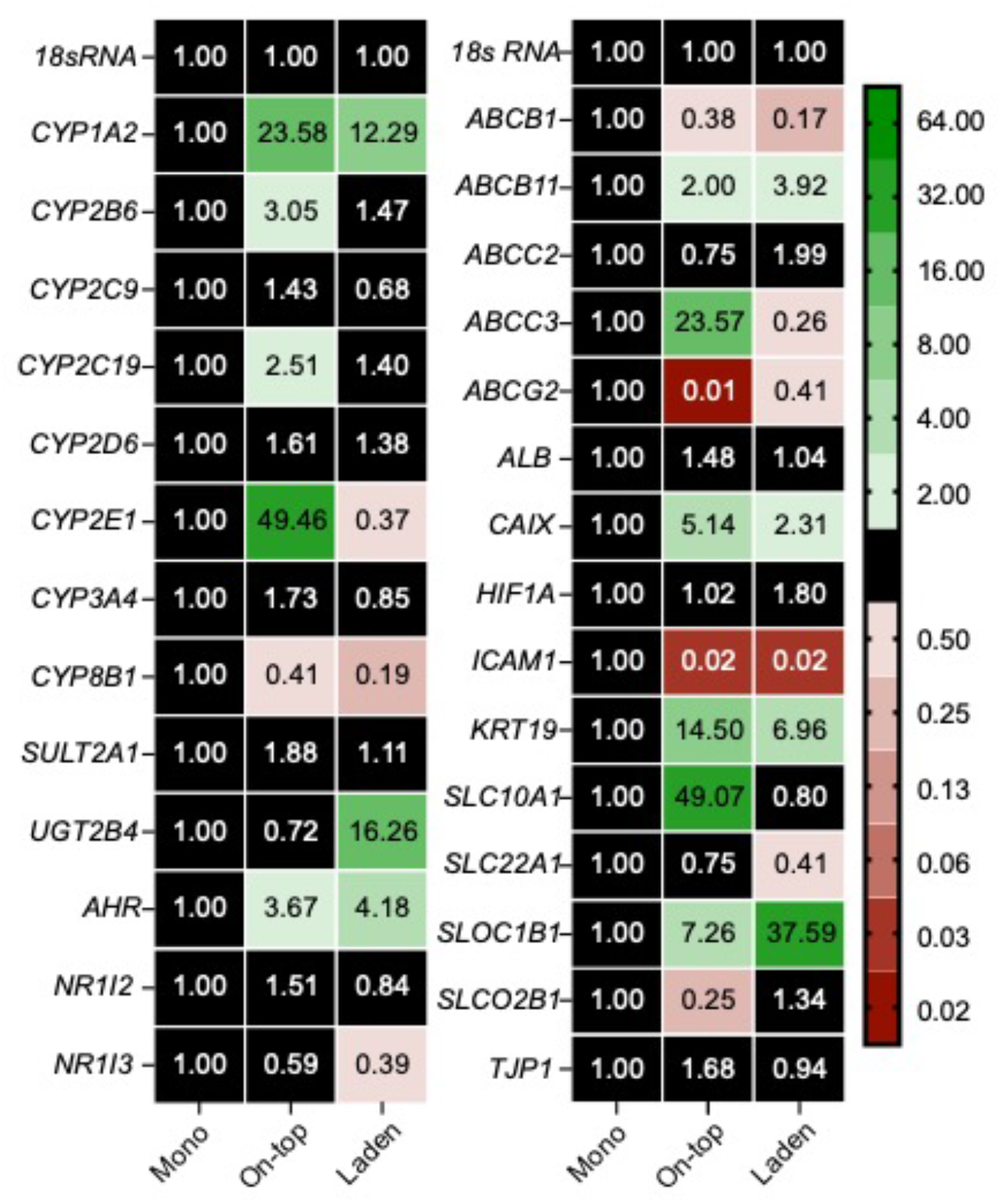
Transcript-level changes of drug-metabolizing enzymes, nuclear receptors, and transport proteins of 1.0×10^6^ HepaRG cells cultured for 8 d under standard culture conditions. Each value is the average fold-change, calculated using the ΔΔCt method, of at least three separate setups prepared from two cryovials of HepaRG cells. NF indicates that no transcript was quantified. A fold-change of greater than 2 or less than 0.50 was considered significant.

### HepaRG polarization and differentiation profiles change when cultured with the paper scaffolds

**Figure 6** also displays transcript-level changes for genes associated with hepatocyte differentiation and polarization. Cytokeratin-19 *(KRT19)* is a cholangiocyte-like marker expressed in hepatocytes forming bile canaliculi.(Higuchi et al. 2014) *KRT19* transcript levels were significantly higher in the on-top (14.5-fold) and laden (6.9-fold) configurations compared to the monolayers. The transcript levels for albumin *(ALB),* a traditional marker for hepatic-like function,(Aleksandr and Müsch 2013) were unchanged between the culture configurations. A previous study found the albumin transcript levels in HepaRG cells cultured as monolayers or suspended in gelatin scaffolds were also unchanged.(Cuvellier et al. 2021)

Transporter proteins are markers of hepatocyte polarization.(Török et al. 2020) We focused on members of the ATP-dependent (ABC) and solute carrier protein families. ABC proteins import nutrients and export xenobiotics.(El-Awady et al. 2017; Hollenstein et al. 2007) The solute carrier (SLC) proteins are a broad class of active transporters that facilitate the movement of small molecule drugs and organic ions (e.g., bile salts) across the cell membrane.(Lin et al. 2015) *ABCB11* levels were significantly increased in the on-top (2.0-fold) and laden (3.9-fold) configurations; *ABCB1* (0.4- and 0.2-fold) and *ABCG2* (0.01- and 0.41-fold) were downregulated in the on-top and laden configurations, respectively. *SLOC1B1* levels increased in the on-top (7.3-fold) and laden (37.6-fold) configurations. *SLC10A1* was unchanged in the on-top configuration but decreased in the laden configuration (0.41-fold).

### Paper scaffold configurations have enhanced post-differentiation responses for drug-metabolizing enzymes compared to monolayers

HepaRG cells in each culture configuration were maintained under standard culture conditions for 6 d, followed by 2 d in a representative PP or PV condition. In the PP condition, cells were maintained at pO_2_ = 11% in basal medium. In the PV conditions, cells were maintained at pO_2_ = 5% in a 1:1 (v/v) ratio of basal culture medium and LWRN-conditioned medium. **Figure 4** shows the urea secretion was unchanged for a given culture condition between the PP and PV conditions: 0.025 mg/mL/h/10^6^ cells in the monolayer; 0.014 mg/mL/h/10^6^ cells in the on-top and laden configurations. The average urea produced in the monolayer configuration was higher than cells in contact with the paper scaffolds in the PP and PV conditions. This decrease in urea production was significant under PP, but not PV conditions. We previously characterized HepaRG monolayers in the same PP and PV culture conditions,(DiProspero et al. 2022) and showed these conditions did not hinder cellular health or viability but did decrease total ATP due to oxygen-mediated glucose metabolism.

**Figure 7** plots drug-metabolizing enzyme activity under PP and PV conditions. Activity increased for CYP1A2, 2D6, 3A4 (probed with midazolam), and UGT enzymes between the PP and PV culture conditions in all three configurations. These increases match the expectations of zonal differences observed in vivo. SULT family activity did not match the expected zonal trend, with increased activity in PV conditions of 8.0-fold in the on-top configuration and 2.4-fold in the laden configuration. In the on-top configuration, enzyme activity between the PP and PV culture conditions increased significantly, 28.7-fold for CYP1A2; 29.9-fold for 2D6; 14.5-fold for 3A4 probed with midazolam and 2.1-fold when probed with testosterone; and 28.6-fold for UGT enzymes. In the laden configuration, the microenvironment changes had less effect than in the on-top configuration, with a 14.4-fold increase in the PV region for CYP1A2, 4.2-fold for 2D6, 4.3-fold (midazolam) and 2.2-fold (testosterone) for 3A4, and 6.7-fold for UGT enzymes.

**Figure. 7.**
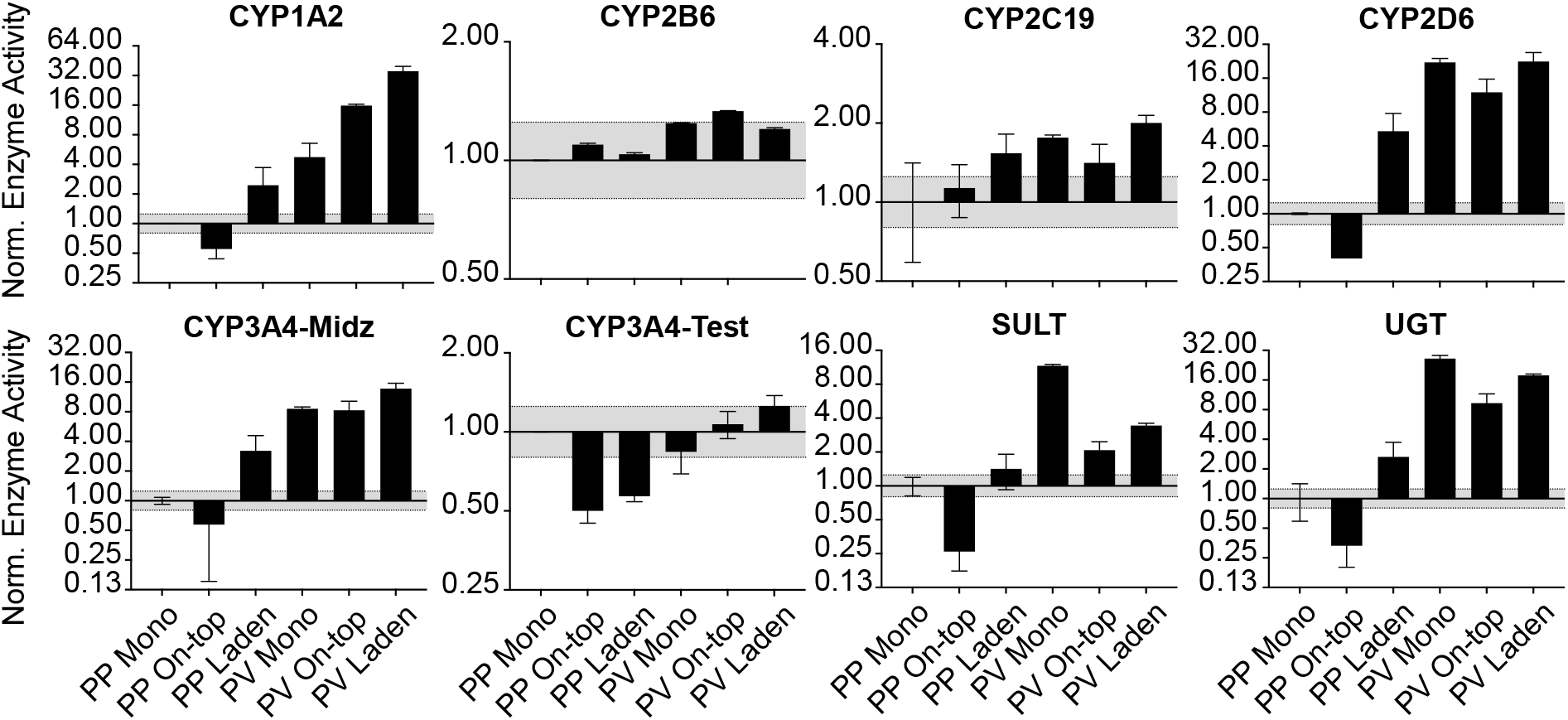
Drug-metabolizing enzyme activity of 1.0×10^6^ HepaRG cells maintained under standard culture conditions for 6 d, followed by 2 d at PP or PV conditions. Activity values were normalized to the monolayer configuration maintained at PP conditions. Each bar represents the average and SEM of two separate setups, prepared from two cryovials of HepaRG cells. The dotted black lines represent a significant difference in activity, based on our LC-MS/MS method.

**Figure 8** displays transcript-level changes in the same cells. Like the activity trends observed in the paper scaffolds, the *CYP, SULT,* and *UGT* enzyme transcripts increased from the PP to PV conditions. In the on-top configuration, there were significant changes in the amount of *CYP1A2* (>64-fold), *2B6* (4.7-fold), *2C9* (20.6-fold), *2C19* (9.3-fold), *3A4* (>64-fold), *UGT2B4* (27.9-fold), *SULT2A1* (>64-fold), and *CYP2D6* (0.2-fold) transcripts. In the laden configuration, changes in most of the transcripts between the PP and PV conditions were less substantial than in the on-top configuration: *CYP1A2* (11.4-fold), *2B6* (11.6-fold), *2C19* (2.9-fold), *3A4* (32.2-fold), *UGT2B4* (54.0-fold), *SULT2A1* (13.8-fold), and *CYP2D6* (0.2-fold). *ALB* transcripts increased in each culture configuration compared to the PP conditions, while the *KTR19* transcripts only increased in the PP on-top configuration.

**Figure. 8.**
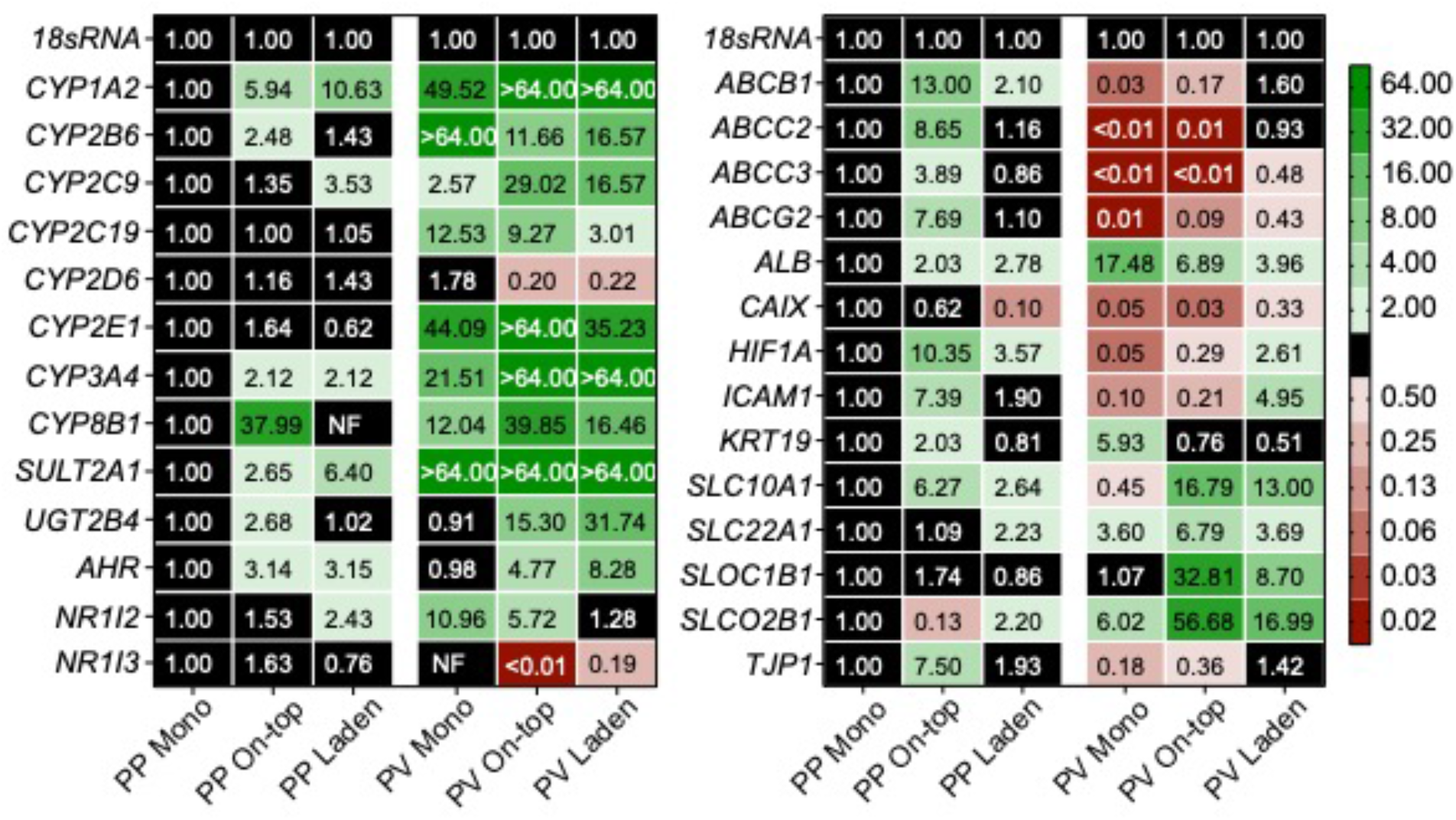
Transcript-level changes of drug-metabolizing enzymes, nuclear receptors, and transport enzymes in HepaRG cells cultured for 6 d under standard conditions, followed by 2 d at either PP or PV conditions. Each value is the average fold-change, calculated using the ΔΔCt method, of at least two separate setups, prepared from two cryovials of HepaRG cells. NF indicates no transcript was quantified. A fold-change >2.0 or < 0.50 was considered significant.

## Discussion

### Paper scaffolds decrease HepaRG drug-metabolizing enzyme activity under standard culture conditions

The paper scaffolds provide a culture region whose volume is defined by a wax-patterned border and the thickness of the paper (40 μm). The exposed cellulose fibers in the unpatterned areas of the scaffold reinforce the thin gels and decrease the chances of cracking or breaking upon physical manipulation. To assess the effect of the paper scaffolds on the drug-metabolizing activity of HepaRG cells, we compared two configurations to a traditional monolayer setup (**Figure 1**). In the on-top configuration, cells were deposited onto a pre-formed collagen gel in a paper scaffold. In the laden configuration, cells were suspended in collagen before seeding into the scaffold. Previous work showed that rat neonatal hepatocytes preferred ECM-containing silk scaffolds with large pore sizes and porosities of 90% void volume.(Janani and Mandal 2021) The sheets of the Whatman 105 paper used in this work have similar void volumes.(Mosadegh et al. 2014) The fluorescent micrographs in **Figure 3** show that the HepaRG cells are evenly distributed across the pre-formed collagen substrates in the monolayer and on-top configurations. The cells in the laden culture concentrate around the paper fibers, forming small aggregates in regions of high fiber density.

The presence of the paper scaffolds does not alter HepaRG viability, with the number of live cells in the on-top and laden configurations within 10% of the monolayer configuration. This decreased viability in the laden configuration could result from pericellular hypoxia in regions of high fiber density, where the small aggregates formed. Previous work showed HepaRG urea production was configuration-dependent, with spheroids secreting 23.5-fold more urea than similar numbers of cells in a sandwich culture.(Li et al. 2019) In our hands, the HepaRG cells produced similar amounts of urea across all three culture configurations, under standard culture conditions. We attribute the differences observed in our paper scaffolds to the absence of large aggregates. However, we hypothesize similar increases in aggregate size and urea production would occur in the laden configuration with higher cell densities.

The drug-metabolizing activity of the paper scaffold-containing cultures was significantly less than the monolayer configurations (**Figure 5**). Cells in the on-top configuration have significantly less CYP1A2, 2D6, 3A4, SULT, and UGT activity. This decrease in activity is not transcriptionally mediated as the cells contained significantly more *CYP1A2, 2B6, 2C19, 2E1,* and *UGT2B4* transcripts in the on-top configuration compared to the monolayer (**Figure 6**). A similar mismatch between transcript and activity occurred between the laden and monolayer culture configurations. While all three culture configurations have equivalent amounts of collagen, non-specific adsorption to the paper fibers in the on-top and laden configurations could sequester the enzyme products. However, a comparison of *ALB* and *KRT19* transcript levels suggest more HepaRG cells adopt a cholangiocyte-like phenotype in the paper scaffolds, a more likely reason for decreased activity than the loss of products to non-specific adsorption on the scaffolds. Janani observed increased CK-19 protein expression for rat hepatocytes maintained on the surface of highly porous silk substrates.(Janani and Mandal 2021) They attributed this post-differentiation change to the increased surface roughness provided by the substrates. The paper fibers may be causing a similar change in the HepaRG cells, a combination of increased cell-cell contact and cell-fiber interactions.

HUES9 human stem cells increase the expression of *ABCC2, ABCB11, ABCG2,* and *SLC10A1* upon completing hepatocyte differentiation and polarization.(Török et al. 2020) We found *ABCB11* increased in the presence of the paper scaffolds. However, *ABCB1* and *ABCG2* levels decreased. We are unaware of a study comparing the transcriptional regulation of HUES9 and HepaRG cells with respect to polarization; such studies will be needed to determine the value of HepaRG transporter expression to model these aspects of PHH post-differentiation. We hypothesize that the cellulose fibers are facilitating polarization. However, further studies will require detailed characterization and high-resolution imaging of cellular morphology, cell-cell contact formation, and the localization of the transport proteins on the cell membrane. Gissen summarized the importance of ECM, cell-surface adherence, and tight junction formation in expressing transport proteins in hepatocyte cells.(Gissen and Arias 2015)

### The paper scaffolds enhanced post-differentiation patterning of CYP activity and transcript in HepaRG cells maintained at physiologic conditions

We systematically compared the effects of physiologic oxygen tensions and the WRN molecules on CYP activity in HepaRG cells maintained as monolayers.(DiProspero et al. 2022) Using a combinatorial experimental design, we showed that physiologic oxygen tension was the primary driver of increased CYP activity between the PP and PV environments. In contrast, the WRN molecules were a minor contributor to their increased activity. The impact of the WRN molecules was confirmed with a comparison to a control treatment, conditioned medium obtained from the parental L-cell line. This previous study guided our experimental design for the current work, allowing us to focus on the dimensionality of the culture and a limited number of culture conditions: a standard condition to compare with other literature studies, a representative PP condition, and a representative PV condition. We confirmed that the paper scaffolds did not deviate from previous results by measuring CYP activity for each culture configuration in the presence of either L- or L-WRN conditioned medium (**Figure S5**). These data show that the WRN molecules increased CYP activity regardless of culture configuration, as in our original study.(DiProspero et al. 2022)

The paper scaffolds enhanced CYP and UGT enzymes compared to the monolayer configuration (**Figure 7**) by contributing to the post-differentiation patterning of the physiologic conditions. Our observations align with zonation and elevated CYP activity in the PV region found in vivo.(Jungermann and Kietzmann 2000) Both the on-top and laden configurations had increased activity of multiple CYPs in the PV conditions. The laden format had a higher increase (7.5-fold) in CYP1A2 activity between the PP and PV conditions than in the on-top configuration (3.4-fold). CYP3A4 activity, when probed with midazolam, was also enhanced by 1.6-fold in the laden configuration compared to the on-top configuration. The activity increases coincide with increased transcripts of *CYP1A2, 2C9, 3A4,* and *UGT2B4.* The paper scaffolds also enhanced other zonal phenotypes compared to the monolayer configuration, including decreased *ALB* and increased *KTR19* transcripts in the PV conditions. Halpern showed that *ALB* is weakly increased in the PP region of the sinusoid,(Halpern et al. 2017) agreeing with our results and suggesting it is zonally regulated. Janani showed hepatocytes maintained at a representative PV oxygen tension had significantly increased levels of *CYP1A2* and *2E1* transcripts and increased expression of CK-19 protein expression compared to the PP region.(Janani and Mandal 2021)

We hypothesize the in vivo-like changes observed in the HepaRG cells between the PP and PV environment are due to increases in the physiological relevance of the culture conditions and configuration. Our previous and current characterization of CYP activity and transcription levels between representative PP and PV conditions highlights the importance of incorporating oxygen partial pressures found in vivo. They also highlight that the WRN molecules have a lesser, albeit important, impact on the drug-metabolizing activity of hepatocytes. Our current work highlighted that the additional context of a tissue-like culture environment further enhances the post-differentiation processing of the HepaRG cells compared to monolayers. Glucose, oxygen, and other signaling molecules are thought to be key regulators of hepatocyte post-differentiation.(Gebhardt et al. 2007; Gebhardt and Hovhannisyan 2010) While further characterization is needed to tease apart the exact mechanism of their role, our results show the paper fibers fine-tune the transcriptional abundance of drug-metabolizing enzymes, transporter proteins, cholangiocyte-like markers, and albumin in HepaRG cells. In agreement with previous studies, our results highlight the potential of the paper-based culture platform to experimentally (and systematically) probe the significance of different microenvironment components on the post-differentiation of hepatocytes.(Agarwal et al. 2020a; Agarwal et al. 2020b; Wang et al. 2018)

## Conclusions

This study compared the effect of both culture dimensionality and physiologically relevant culture conditions on the activity and transcript levels of drug-metabolizing enzymes in HepaRG cells. To make these comparisons, we exposed three different culture configurations to standard culture conditions, a representative PP oxygen tension and a representative PV oxygen tension with the addition of WRN molecules. In the three culture configurations, cells were maintained as a monolayer on a collagen slab (monolayer configuration), placed on top of a collagen-containing paper scaffold (on-top configuration), or mixed into collagen gel and then seeded into a paper-scaffold (laden configuration). Under standard conditions, the presence of the paper scaffolds decreased the overall CYP activity of the HepaRG cells compared to the monolayer cultures. Transcript analysis showed the paper scaffolds increased cellular polarity and the number of cholangiocyte-like cells. When comparing HepaRG maintained under the PP and PV culture conditions, we observed increased drug-metabolizing enzyme activity and transcripts in the culture configurations containing the paper scaffolds. Transcripts highlight the paper scaffold’s role in increasing the polarization of the cells compared to cells on collagen-only monolayers.

Our results also highlight the importance of context when designing cell-based assays. Incorporating a tissue-like environment and physiologically relevant oxygen tensions highlights the need to reconsider the status quo of monolayer cultures exposed to atmospheric oxygen tensions when trying to predict or extrapolate hepatocyte responses or small molecule toxicities in vivo. Our results also highlight the utility of the paper-based culture platform for continued studies, as the cellulose fibers do not decrease the viability, health, or post-differentiation patterning of the HepaRG cells under physiologically relevant conditions.

## Acknowledgements

This work was supported by the National Institute of General Medical Sciences (NIGMS) through Grant Number R35GM128697. We thank the University of North Carolina’s Department of Chemistry Mass Spectrometry Core Laboratory, especially Dr. Brandie Ehrmann, for their assistance with mass spectrometry analysis. A portion of this work was performed on Core instrumentation that was funded in part by the University of North Carolina’s School of Medicine Office of Research. We also thank Mr. Tyler Larson for their assistance with instrument troubleshooting and helpful discussions as we prepared this manuscript.

## Author Contribution

Thomas J. DiProspero and Matthew R. Lockett contributed to the study conception and design. All authors contributed to material preparation, data collection, and analysis. The first draft of the manuscript was written by Thomas J. DiProspero, and all authors commented on previous versions of the manuscript. All authors read and approved the final manuscript.

## Declarations

### Competing Interests

The authors have no relevant financial or non-financial interests to disclose.

### Ethical Approval

Not Applicable. This work did not involve animal, human, or clinical samples.

### Funding

This work was funded by the National Institute of General Medical Sciences (NIGMS) through Grant Number R35GM128697.

### Availability of Data

Additional data and information can be found in the Supporting Information. The raw values and worked-up datasets are available for further analyses, please contact the corresponding author.

## Notes

### Competing Interest Statement

The authors have declared no competing interest.

