## Supporting Information for "HepaRG cells undergo increased levels of post-differentiation patterning in physiologic conditions when maintained as 3D cultures in paper-based scaffolds"

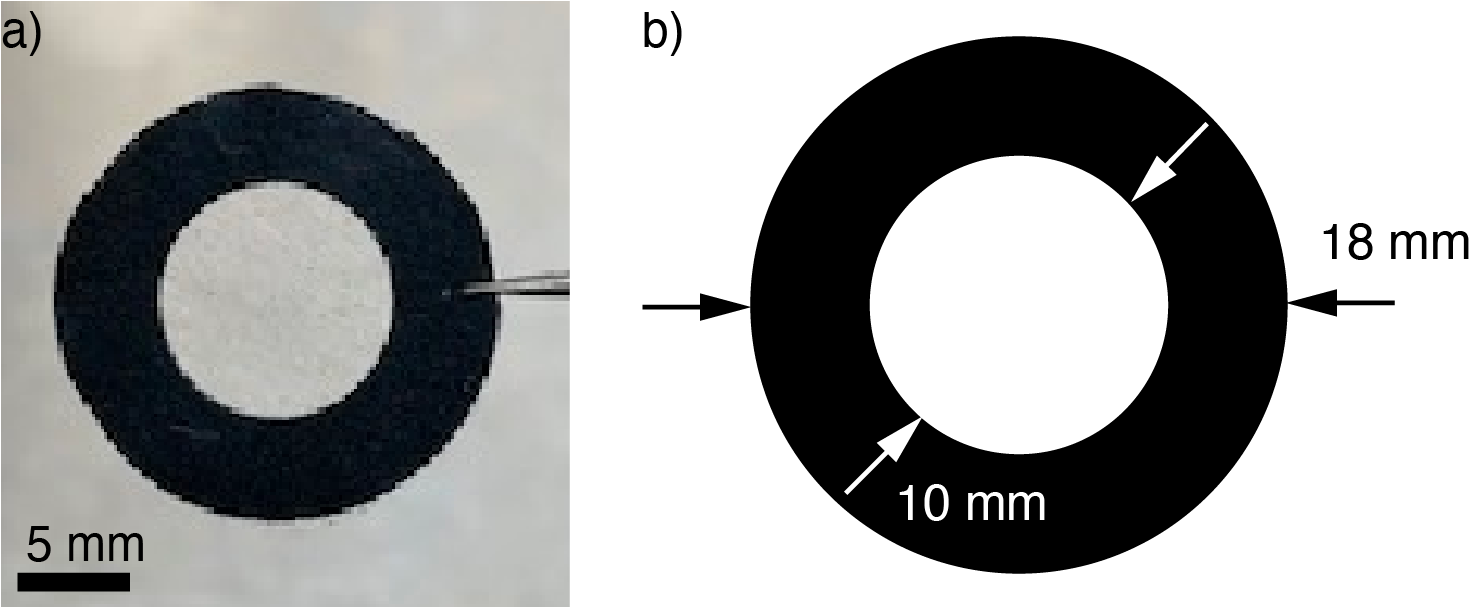

**Figure S1.** Picture (a) and schematic (b) of paper scaffolds used in this work. Patterns were designed with Adobe Illustrator and patterned onto Whatman 105 lens paper using a Xerox ColorQube 8570 wax printer.

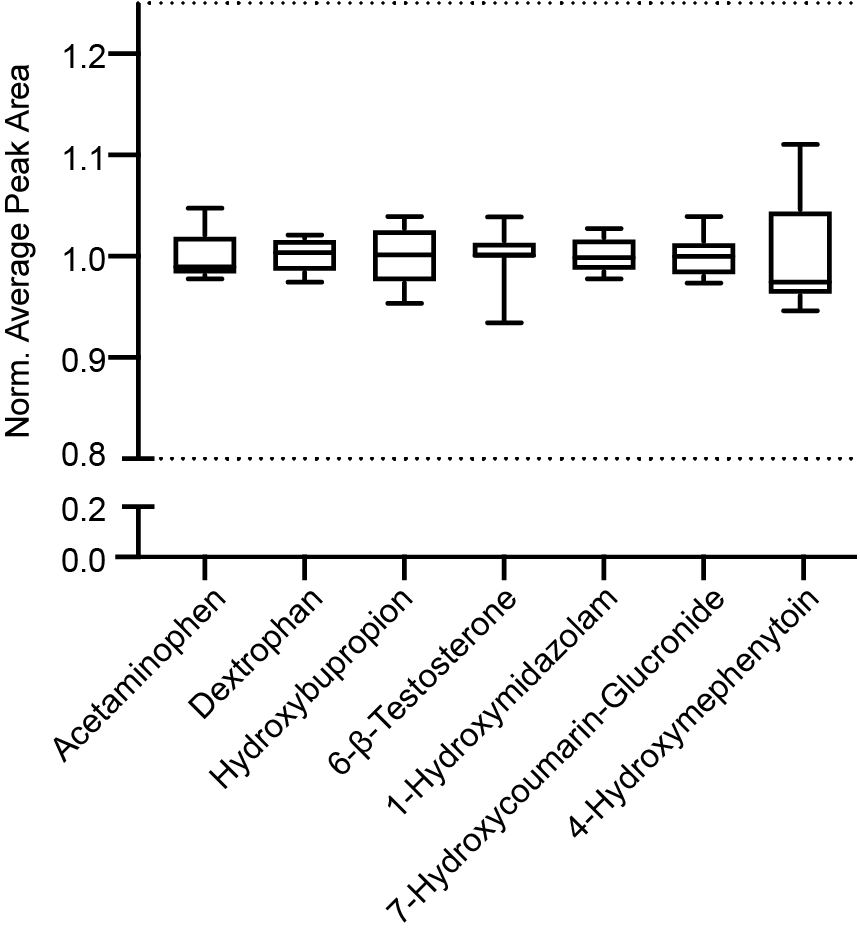

**Figure S2.** Repeated injections of acetaminophen and six CYP and UGT reaction products, quantified with the LC-MS/MS method outlined in the manuscript. The product mixture was prepared in Optima water with a final concentration of 10 μM of each analyte. In this plot, the whiskers are the range; the upper box is the 75^th^ percentile; the lower box is the 25^th^ percentile; and the line represents the median measurement. The dashed lines represent four standard deviations from the mean, accounting for over 99.99% of the deviation on a normal bell curve.

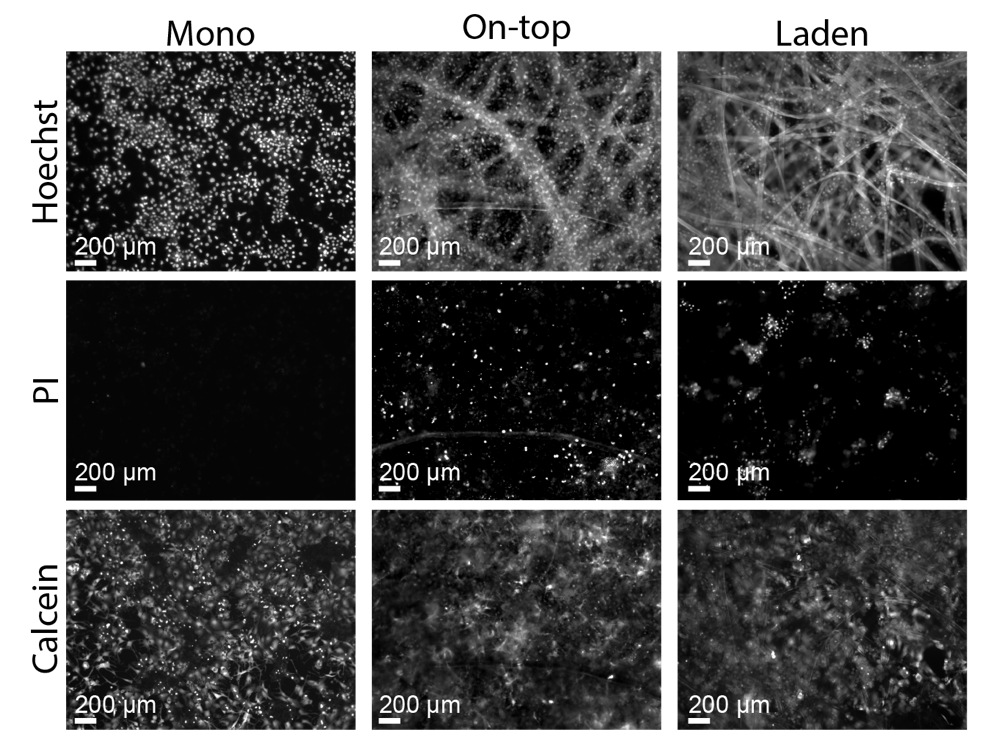

**Figure S3.** Representative images of 1.0x10^6^ HepaRG cells stained with Hoechst 33343, propidium iodide (PI) and calcein-AM in the (left) monolayer, (middle) on-top, or (right) laden culture configurations.

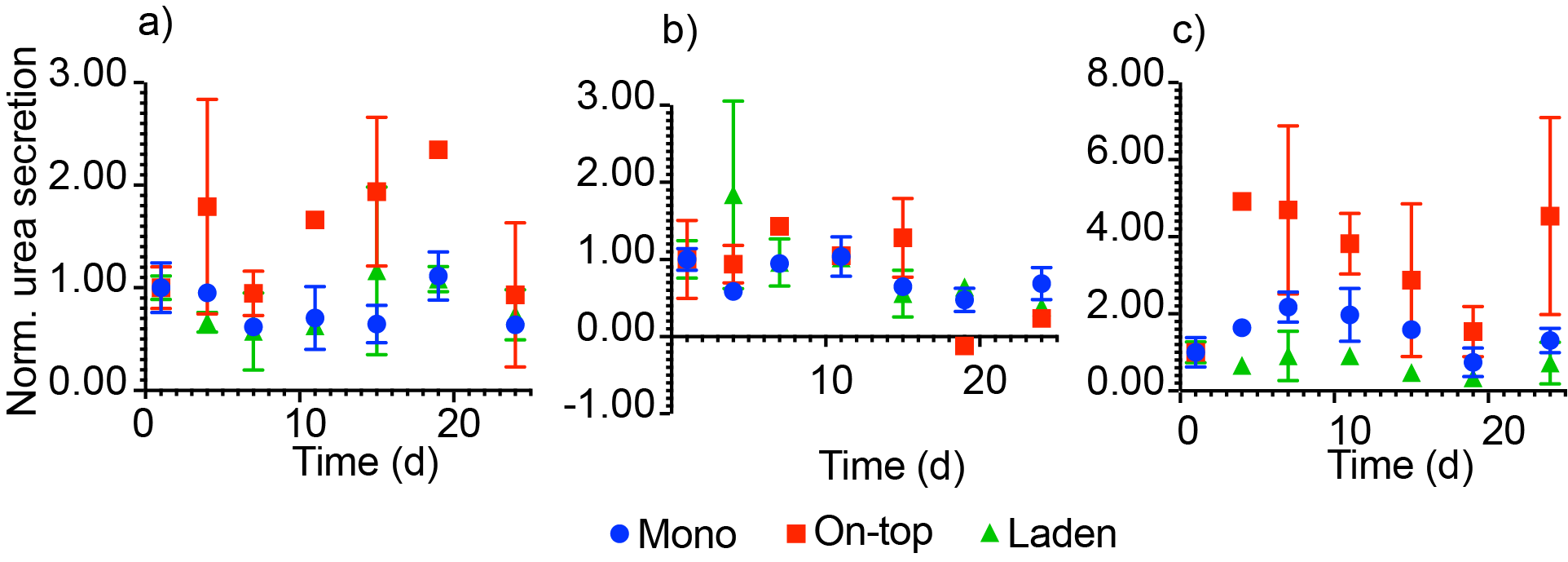

**Figure S4.** Urea secretion of HepaRG cells for 24 d after deposition, maintained in (a) standard, (b) L-Cell, or (c) L-WRN treated medium for cells seeded in the monolayer, on-top, and laden culture configurations. Values are normalized to the urea produced 1 day after seeding. Urea was quantified directly from culture medium samples using a colorimetric assay. Each point represents the average value from two separately prepared setups; the bars represent the range of those values.

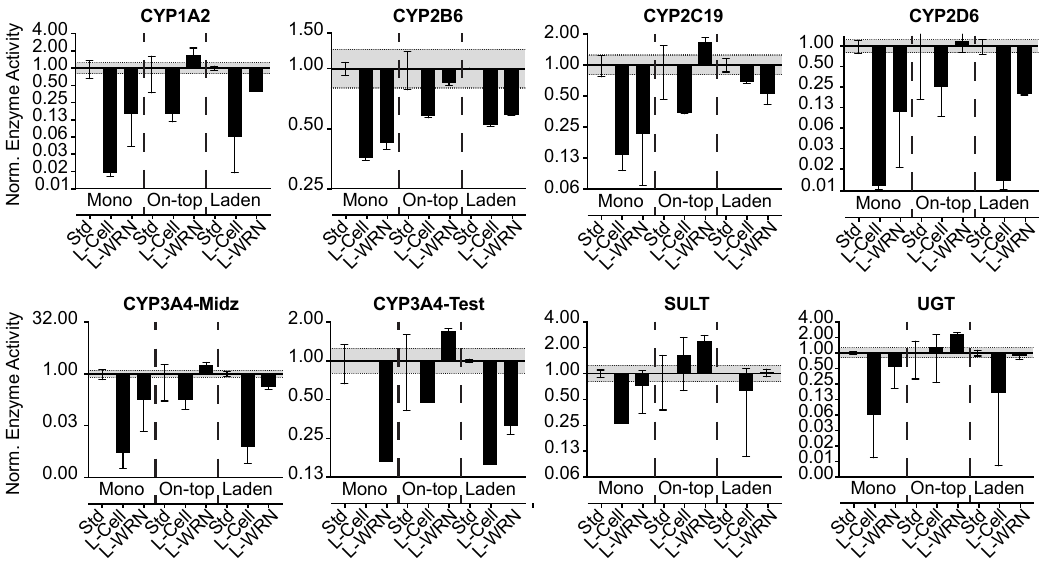
**Figure S5.** Drug-metabolizing enzyme activity of 1.0x10^6^ HepaRG cells maintained under standard culture conditions for 6 d followed by 2 d with standard, L-cell, or L-WRN conditioned medium. Activity values were normalized to the standard medium treatment in each configuration. Each bar represents the average and SEM of three separate setups, prepared from two cryovials of HepaRG cells. The dotted black lines represent a significant difference in activity, based on the LC-MS/MS method used.

**Table S1**. Selective reaction monitoring (SRM) MS/MS transitions for each enzyme product.

| **Enzyme** | **Substrate** | **Final concentration (μM)** | **Product/Isotope standard** | **Declustering voltage (V)** | | **Ion mode** | | **Parent ion (m/z)** | | **Product ion (m/z)** | | **Collision energy (eV)** |
| --- | --- | --- | --- | --- | --- | --- | --- | --- | --- | --- | --- | --- |
| CYP1A2 | Phenacetin | 100 | Acetaminophen | 6 | Positive | | 152.2 | | 110.0 | | 15 | |
|  |  |  |  |  |  |  | 152.2 | | 65.0 | | 30 | |
| CYP2B6 | Bupropion | 50 | Hydroxybupropion | 6 | Positive | | 256.0 | | 238.1 | | 8 | |
|  |  |  |  |  |  |  | 256.0 | | 130.1 | | 47 | |
| CYP2C19 | (*S*)-mephenytoin | 100 | 4-hydroxymephenytoin | 6 | Negative | | 232.9 | | 190.1 | | 19 | |
|  |  |  |  |  |  |  | 232.9 | | 161.0 | | 25 | |
| CYP2D6 | Dextromethorphan | 100 | Dextrorphan | 4 | Positive | | 258.0 | | 157.1 | | 36 | |
|  |  |  |  |  |  |  | 258.0 | | 199.1 | | 25 | |
| CYP3A4 | Midazolam | 5 | 1-hydroxymidazolam | 6 | Positive | | 342.0 | | 324.1 | | 19 | |
|  |  |  |  |  |  |  | 342.0 | | 168.1 | | 36 | |
| CYP3A4 | Testosterone | 50 | 6-β-testosterone | 8 | Positive | | 305.2 | | 269.2 | | 13 | |
|  |  |  |  |  |  |  | 305.2 | | 105.1 | | 36 | |
| SULT | 7-hydroxycoumarin | 100 | 7-hydroxycoumarin sulfate | 6 | Negative | | 240.7 | | 161.0 | | 20 | |
|  |  |  |  |  |  |  | 240.7 | | 133.0 | | 34 | |
| UGT | 7-hydroxycoumarin | 100 | 7-hydroxycoumarin glucuronide | 6 | Negative | | 337.0 | | 161.0 | | 29 | |
|  |  |  |  |  |  |  | 337.0 | | 175.0 | | 13 | |
| CYP1A2 |  |  | Acetaminophen-d_4_ | 5 | Positive | | 152.2 | | 110.0 | | 15 | |
|  |  |  |  |  |  |  | 152.2 | | 65.0 | | 30 | |
| CYP2B6 |  |  | Hydroxybupropion-d_6_ | 6 | Positive | | 262.0 | | 244.1 | | 12 | |
|  |  |  |  |  |  |  | 262.0 | | 130.0 | | 43 | |
| CYP2C19 |  |  | 4’-hydroxymephenytoin-d_3_ | 6 | Negative | | 236.0 | | 193.1 | | 17 | |
| CYP2D6 |  |  | Dextrorphan-d_3_ | 4 | Positive | | 261.2 | | 157.1 | | 40 | |
|  |  |  |  |  |  |  | 261.2 | | 199.1 | | 25 | |
| CYP3A4 |  |  | 1-hydroxymidazolam-[^13^C_3_] | 10 | Positive | | 345.1 | | 327.1 | | 18 | |
|  |  |  |  |  |  |  | 345.1 | | 171.1 | | 36 | |
| CYP3A4 |  |  | 6-β-testosterone-d_7_ | 5 | Positive | | 312.2 | | 276.2 | | 19 | |
|  |  |  |  |  |  |  | 312.2 | | 294.1 | | 11 | |
| SULT |  |  | 7-hydroxycoumarin sulfate-d_5_ | 0 | Negative | | 245.1 | | 165.0 | | 31 | |
|  |  |  |  |  |  |  | 245.1 | | 137.1 | | 39 | |
| UGT |  |  | 7-hydroxycoumarin-^13^C_6_-glucuronide | 16 | Negative | | 342.9 | | 167.1 | | 32 | |
|  |  |  |  |  |  |  | 342.9 | | 139.1 | | 29 | |

**Table S2**. qPCR primers for each gene evaluated.

| **Gene Symbol** | **Protein Abbreviation** | **Main Function** | **Forward Primer (5’ – 3’)** | **Reverse Primer (5’ – 3’)** | **Efficiency (%)** |
| --- | --- | --- | --- | --- | --- |
| *18sRNA* | 18s rRNA | Ribosome | CGCCGCTAGAGGTGAAATTC | TTGGCAAATGCTTTCGCTC | 107.5 |
| *CYP1A2* | CYP1A2 | Phase I Enzyme | CTTCGGACAGCACTTCCCTG | AGGGTTAGGCAGGTAGCGAA | 103.9 |
| *CYP2B6* | CYP2B6 | Phase I Enzyme | TTCCTACTGCTTCCGTCTATC | GTGCAGAATCCCACAGCTCA | 101.4 |
| *CYP2C9* | CYP2C9 | Phase I Enzyme | TCCCTGACTTCTGTGCTACATG | ACTGGAGTGGTGTCAAGGTTC | 113.9 |
| *CYP2C19* | CYP2C19 | Phase I Enzyme | CAACAACCCTCGGGACTTTA | GTCTCTGTCCCAGCTCCAAG | 106.2 |
| *CYP2D6* | CYP2D6 | Phase I Enzyme | ACCAGGCTCACATGCCCTA | TTCGATGTCACGGGATGTCAT | 103.7 |
| *CYP2E1* | CYP2E1 | Phase I Enzyme | TTGAAGCCTCTCGTTGACCC | CGTGGTGGGATACAGCCAA | 109.9 |
| *CYP3A4* | CYP3A4 | Phase I Enzyme | TCACAAACCGGAGGCCTTTT | TGGTGAAGGTTGGAGACAGC | 100.4 |
| *CYP8B1* | CYP8B1 | Phase I Enzyme | TGCACATGGACCCTGACATC | GTGTCAGGGTCCACCAACTC | 91.9 |
| *SULT2A1* | SULT2A1 | Phase II Enzyme | TGAGGAGCTGAAACAGGACAC | AAGTCTTCAGCTTGGGCCAC | 106.6 |
| *UGT2B4* | UGT2B4 | Phase II Enzyme | ACACATGAAGGCCAAGGGAG | GAACCAGGTGAGGTCGTGG | 94.3 |
| *AHR* | AhR | Transcription Factor | CTTCCAAGCGGCATAGAGAC | AGTTATCCTGGCCTCCGTTT | 101.5 |
| *NR1I2* | PxR | Transcription Factor | CCAGGACATACACCCCTTTG | CTACCTGTGATGCCGAACAA | 104.3 |
| *NR1I3* | CaR | Transcription Factor | TGATCAGCTGCAAGAGGAGA | AGGCCTAGCAACTTCGCATA | 102.6 |
| *ABCB1* | P-gp | Efflux Pump | GCCAAAGCCAAAATATCAGC | TTCCAATGTGTTCGGCATTA | 93.6 |
| *ABCB11* | BSEP | Bile Salt Exporter Pump | TGATCCTGATCAAGGGAAGG | TGGTTCCTGGGAAACAATTC | 102.5 |
| *ABCC2* | MRP2 | Transporter (Excretion) | TGAGCAAGTTTGAAACGCACAT | AGCTCTTCTCCTGCCGTCTCT | 99.6 |
| *ABCC3* | MRP3 | Transporter (Excretion) | GTCCGCAGAATGGACTTGAT | TCACCACTTGGGGATCATTT | 108.5 |
| *ABCG2* | BCRP | Transporter (Excretion) | TGCAACATGTACTGGCGAAGA | TCTTCCACAAGCCCCAGG | 101.5 |
| *SLC10A1* | NTCP | Transporter (Uptake) | GGGACATGAACCTCAGCATT | CGTTTGGATTTGAGGACGAT | 101.4 |
| *SLC22A1* | OCT1 | Transporter (Uptake) | TAATGGACCACATCGCTCAA | AGCCCCTGATAGAGCACAGA | 104.5 |
| *SLOC1B1* | OATP1B1 | Transporter (Uptake) | GCCCAAGAGATGATGCTTGT | ATTGAGTGGAAACCCAGTGC | 97.3 |
| *SLCO2B1* | OATP2B1 | Transporter (Uptake) | TGATTGGCTATGGGGCTATC | CATATCCTCAGGGCTGGTGT | 106.5 |
| *ALB* | Albumin | Globular Protein | TGAGCAGCTTGGAGAGTACA | GTTCAGGACCACGGATAGAT | 124.1 |
| *CAIX* | Carbonic Anhydrase 9 | Hypoxia Marker/ Hydration of CO_2_ | GGGCCCGGAAGAAAACAGT | TCTTCCAAGCGAGACAGCAAC | 104.8 |
| *HIF1* | HIF1α | Hypoxia response | CATAAAGTCTGCAACATGGAAGGT | ATTTGATGGGTGAGGAATGGGTT | 100.9 |
| *KRT19* | CK-19 | Biliary-like/Progenitor  Cell Marker | TTTGAGACGGAACAGGCTCT | AATCCACCTCCACACTGACC | 100.8 |
| *TJP1* | ZO-1 | Tight-Junction | CGAGTTGCAATGGTTAACGGA | TCAGGATCAGGACGACTTACTGG | 106.9 |
